# Dynamics of efficient ensemble coding

**DOI:** 10.1101/2024.12.20.629841

**Authors:** Long Ni, Alan A. Stocker

## Abstract

Ensemble coding creates compressed representations of a stimulus array. When discriminating the ensemble average against a reference, however, items in the ensemble with feature values closer to the reference are typically weighed stronger. We have recently shown that this “robust averaging” behavior can be explained as a form of efficient coding, where the sensory encoding precision is dynamically adapted to efficiently represent ensemble stimuli according to their overall distribution relative to a trial-by-trial varying reference. However, the specific mechanisms underlying such dynamic, efficient ensemble coding remain unknown. Here, we demonstrate that the relative timing between the presentation of the reference and the stimulus ensemble strongly affects efficient ensemble coding. We systematically probed participants’ decision behavior for varying time intervals between the presentation of the ensemble and the reference. We found that efficient ensemble coding was only clearly established when reference and ensemble were simultaneously presented. It was much weaker when the ensemble preceded the reference, and largely absent when the ensemble followed the reference. As captured by our model, reduced efficient ensemble coding thereby coincided with decreased decision accuracy in those asynchronous conditions. Our results indicate that any temporal offset between the ensemble and reference stimuli substantially disrupts the dynamic and efficient allocation of coding resource. This suggests that efficient ensemble coding is not the result of a preparatory attentional process nor due to evidence selection at the decision stage. Rather, it arises from a fast interaction between the simultaneously evoked, sensory representations of reference and ensemble stimuli.

## Introduction

Human vision is constrained by its limited information processing capacity. To alleviate this limitation, the visual system often establishes rapid summary representations of complex visual displays, a process known as ensemble coding. Ensemble coding helps the system mitigate its information processing bottleneck and achieve a stable and consistent representation of the environment given its inherently noisy representations of individual stimuli.

Numerous studies have demonstrated that observers can quickly and effortlessly extract the average value of both low- and high-level visual features of a stimulus ensemble (for a review, see Whitney and Yamanashi-Leib (2018)). However, not every item in the ensemble contributes equally to the estimated ensemble average. Typically, observers tend to overweigh stimuli that are attended to (De Fockert and Marchant, 2008; Choi and Chong, 2020), are more salient (Im et al., 2015; Kanaya et al., 2018; Iakovlev and Utochkin, 2021), and are displayed in the left or central visual field (Li and Yeh, 2017; Pascucci et al., 2021; Tiurina et al., 2024; Dandan et al., 2023). Furthermore, when discriminating the ensemble average against a reference, items with feature values similar to the reference (inliers) contribute stronger to the decision compared to those with more dissimilar feature values (outliers) (de Gardelle and Summerfield, 2011; Li et al., 2017; Vandormael et al., 2017). Such “robust averaging” has been considered evidence that human observers are not optimally integrating sensory information across space (de Gardelle and Summerfield, 2011; Rahnev and Denison, 2018).

However, we have previously demonstrated that robust averaging naturally emerges from an optimal integration process that is constrained by limited coding resources (Ni and Stocker, 2023). We introduced a hierarchical Bayesian decision model whose sensory likelihood functions are shaped by the assumption that the visual system rapidly forms efficient sensory representations according to the overall statistics of the ensemble stimuli in the experiment. We showed that this “efficient ensemble coding” model accurately accounts for robust averaging behavior reported in multiple ensemble decision-making studies.

An important aspect is that the ensemble statistics are measured *relative* to the reference in the decision task. Not only does this imply that the visual system can successfully learn the statistics of a relative stimulus distribution over the course of the experiment. But more importantly, because the reference changes trial by trial, it suggests that the visual system can rapidly deploy the corresponding efficient encoding kernel for the specific reference value in each trial. While traditionally, efficient coding models in visual perception have been mainly considered with regard to the long-term, absolute statistical distributions in the visual diet (e.g., Wei and Stocker, 2015; Ganguli and Simoncelli, 2014; Polania et al., 2019; Zhang and Stocker, 2022), more recent results suggest that efficient codes can dynamically operate at shorter timescales relevant for perceptual learning or adaptation (Schaffner et al., 2023; Mao et al., 2024) or even faster, during single stimulus exposures (Zhang et al., 2024). Thus, fast, efficient ensemble coding is plausible. Yet the underlying mechanism of this dynamic coding process remains unknown, and is the subject of the present study.

Here, we examined how the relative timing between the presentation of the reference and the ensemble stimuli affects efficient ensemble coding. We addressed this question by systematically varying the inter-stimulus interval (ISI) between the ensemble and the reference stimuli in an ensemble discrimination task. We found that efficient ensemble coding (i.e., robust averaging) and associated high decision accuracy only occurred when the ensemble and the reference stimuli were simultaneously presented. Efficient ensemble coding and decision performance dropped markedly when presentation of the ensemble either preceded the reference stimulus, or vice versa. Our efficient ensemble coding model allowed us to precisely quantify these effects as a function of ISI duration, revealing that a simultaneous access of the ensemble and reference stimuli is crucial for the dynamic allocation of encoding resource in ensemble coding on a trial-by-trial basis. Our results suggest that efficient ensemble coding is quickly established via interactions of evoked sensory responses representing the stimulus ensemble and the reference during a single trial.

## Methods

### Experiment

Nine subjects (five women, four men, age 18 to 21) participated in the experiment. All subjects had normal or corrected-to-normal vision. Each subject received course credits for their participation. The study adhered to the Declaration of Helsinki and was approved by the Institutional Review Board of the University of Pennsylvania.

### Apparatus and stimuli

Stimuli were displayed on a special purpose monitor (VIEWPixx3D) with a resolution of 1920 × 1080 pixels and a refresh rate of 120 Hz. Subjects were seated in a darkened room and viewed the display from a distance of 83.5 cm, with their head resting on a chin rest. All stimuli were generated with the Psychophysics Toolbox 3 in MATLAB (Brainard and Vision, 1997). The background was medium gray with luminance of 40 *cd/m*^2^. Each ensemble stimulus consisted of 12 individually oriented gratings in circular apertures, each aperture spanning 1.2 deg of visual angle (0.5 Michelson Contrast). A reference grating was presented at the center of the screen within an aperture spanning 1 deg of visual angle. The 12 ensemble gratings were equally spaced along a virtual circle centered at the center of the screen with a radius of 4.05 deg visual angle. On each trial, the positions of the 12 ensemble gratings were concurrently shifted along the circle by a random angle. Ensemble and reference gratings were identical in spatial frequency (2.17 cycles/deg) with randomized phases in each trial.

### Trial procedure

A trial started with a white fixation point at the center of the screen, presented for 500 ms. An ensemble (or reference) stimulus was then presented for 500 ms. After a variable inter-stimulus interval (ISI) randomly selected between 0 and 900 ms, the reference (or, alternatively, the ensemble) was presented for another 500 ms. In 1/5 of the trials, the reference and ensemble stimuli were displayed simultaneously for 500 ms. Upon seeing a response cue, the subject was instructed to discriminate the generative mean orientation of the ensemble against the reference orientation and to press ‘F’ on the keyboard for counter-clockwise (ccw) and ‘J’ for clockwise (cw) choices, respectively. After each response, auditory feedback indicated a correct (high-pitched tone) or an incorrect decision (low-pitched tone). Feedback was given to provide subjects with information necessary to accurately learn the ensemble distribution and the task. Figure 1 illustrates the overall trial design and the three different trial conditions.

**Figure 1:**
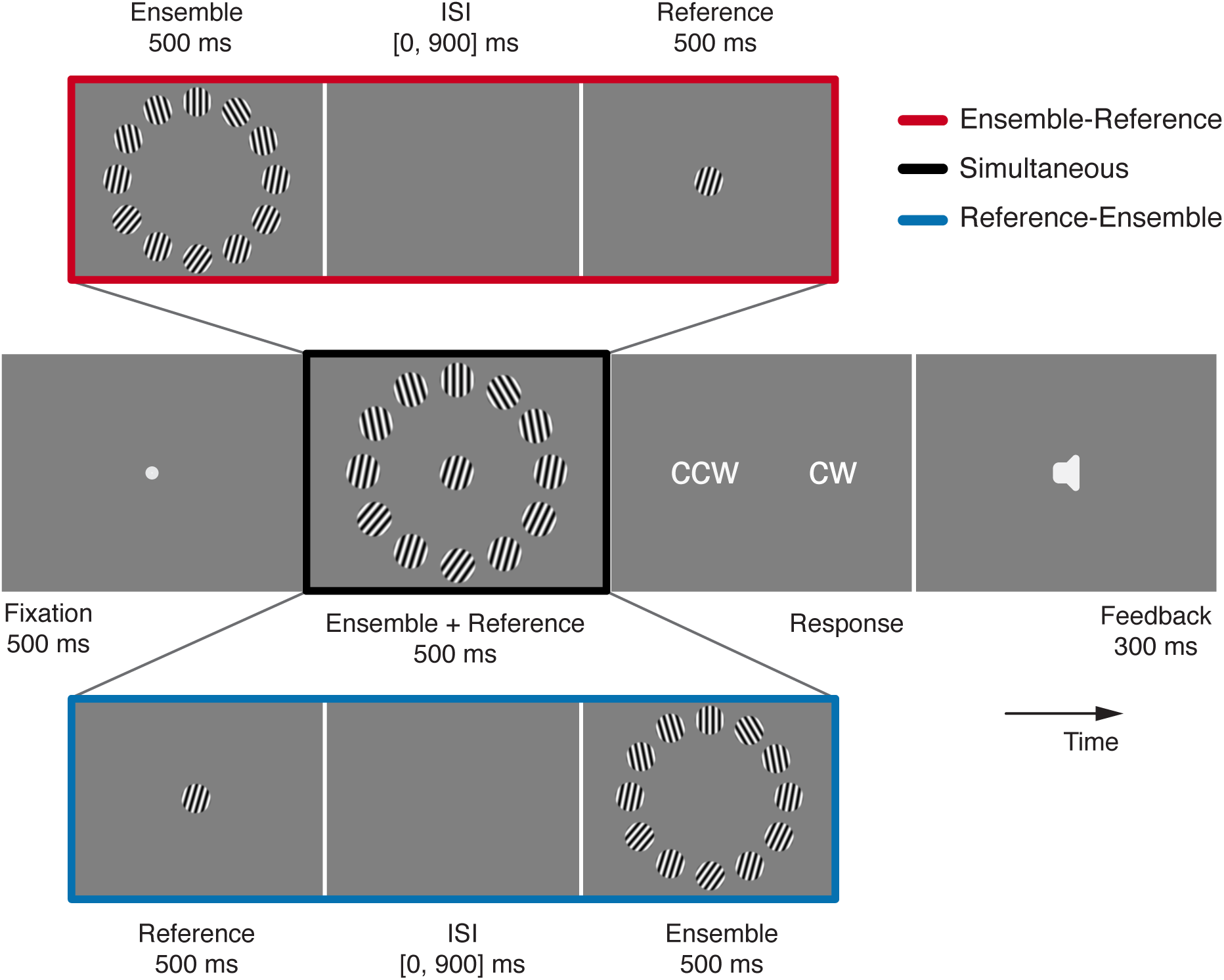
Ensemble discrimination task. A trial started with a fixation dot displayed on center for 500 ms, followed by a reference grating (or ensemble gratings) for 500 ms. After a variable inter-stimulus interval (ISI) randomly selected between 0 and 900 ms, the ensemble (or reference) was presented for another 500 ms. In 1/5 of the trials (randomly selected), the ensemble and reference stimuli were displayed simultaneously (black). Upon viewing the response cue, subjects judged whether the orientation of the reference was clockwise (cw) or counter-clockwise (ccw) relative to the (generative) mean orientation of the ensemble gratings. Subjects received auditory feedback about their choice at the end of each trial.

On each trial, the ensemble orientations were sampled from a Gaussian distribution with a standard deviation of 10 deg and a mean randomly selected from a predefined set {±5, ±10 and ±20} deg relative to the reference orientation (Fig. 2a). Taken together, this created an approximately Gaussian overall distribution of ensemble orientations centered at the reference (Fig. 2b, left panel). The reference orientation itself was randomly sampled for each trial from the range [-90, 90] deg.

**Figure 2:**
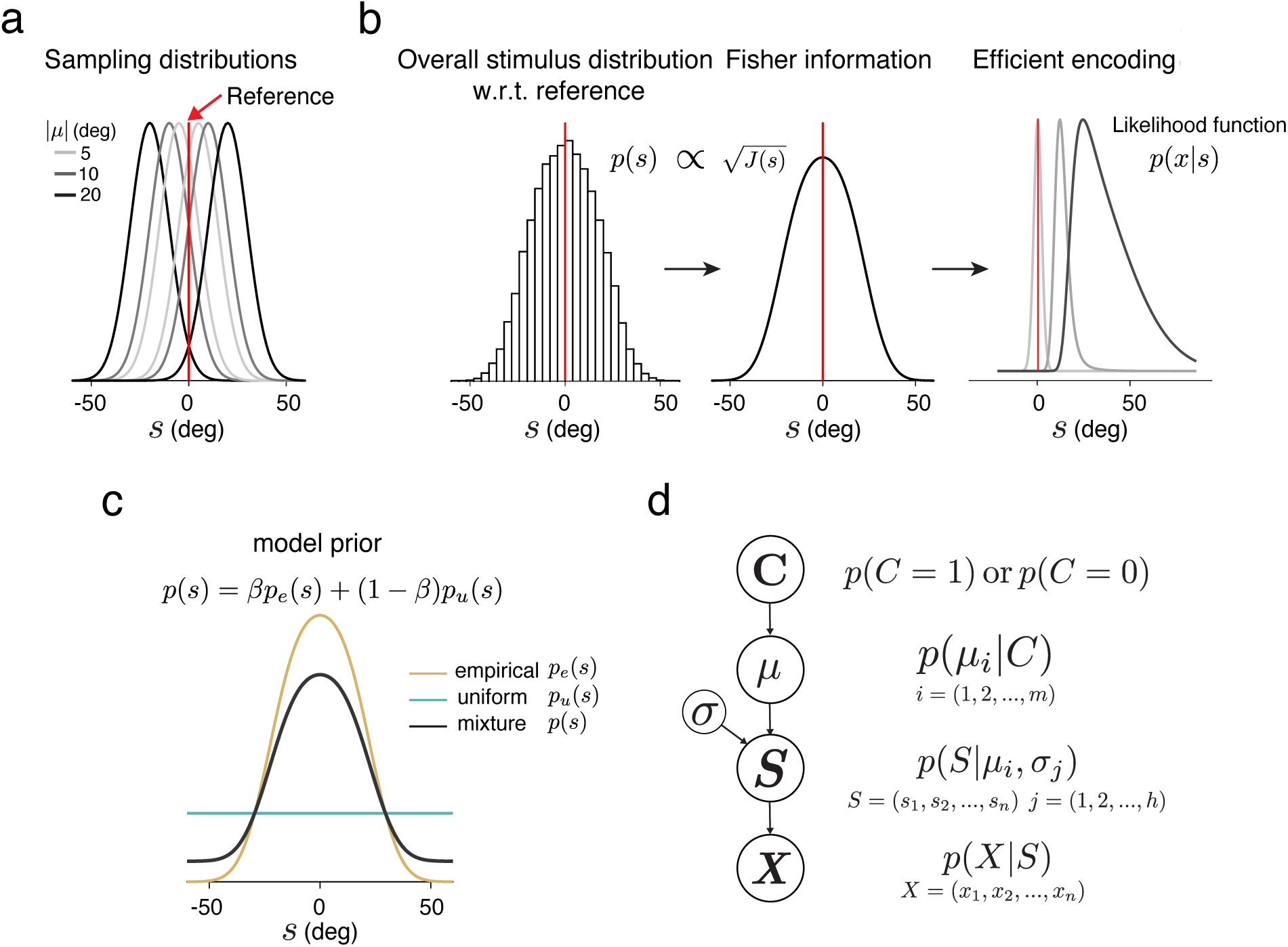
Efficient ensemble coding model (Ni and Stocker, 2023). (a) On each trial, orientations of the ensemble items were randomly sampled from a Gaussian distribution with a fixed variance and a mean randomly selected from a predefined set relative to the reference (red vertical line). (b) Histogram showing the overall orientation distribution of the ensemble stimuli relative to the reference (left panel). Efficient sensory encoding predicts that the encoding precision of stimulus orientation, measured in Fisher information *J* (*s*), is proportional to this overall orientation distribution *p*(*s*) (prior). The inhomogeneous sensory encoding results in likelihood functions *p*(*x*|*s*) that are narrow and symmetric for sensory measurements *x* close to the reference (red vertical line) and wider and asymmetric as *x* is further away (right panel). (c) For the model, we assumed that the stimulus prior *p*(*s*) is a weighted mixture of the overall stimulus distribution in the experiment *p_e_*(*s*) (black line), and a uniform distribution *p_u_*(*s*) (blue line). Free parameter *β* characterizes how much the observer’s sensory encoding is adapted to the overall stimulus distribution. (d) Hierarchical, generative model of the ensemble discrimination task. On each trial, the orientations *S* of the items in the stimulus ensemble represent samples from a Gaussian distribution with varying means *µ* depending on the category of the stimulus ensemble *C* (i.e., which side of the category boundary *µ* is). An observer has to infer the correct category *C* based on noisy observations *X* = (*x*_1_*, …, x*_12_) of each item’s orientation.

### Experimental procedure

Each subject completed 480 simultaneous, 960 ensemble-reference, and 960 reference-ensemble trials. The resulting 2400 trials were randomly interleaved and equally divided into four sessions.

Each subject maximally performed two sessions per day with a 15-min break in-between. Within each session, there were breaks after every 100 trials. Each session took approximately half an hour to complete. Before data collection, subjects underwent extensive training with simultaneous trials to ensure they had been well adapted to the overall ensemble distribution.

### Analysis and model

#### Regression analysis

We extracted the weight with which each item in the stimulus ensemble contributed to subjects’ choices using logistic regression. The regression model assumes subjects’ decision probability *p*(*Ĉ* = {cw, ccw}) (i.e., probability to perceive the ensemble orientation average clockwise/counterclockwise of the reference orientation) is described by

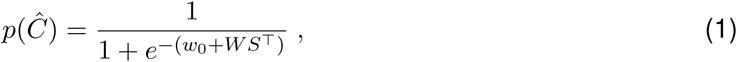

where 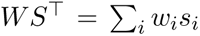, with *w_i_* representing the weight and *s_i_* the orientation of each item *i* in the ensemble, and *w*_0_ an offset parameter. The position *i* of each item in the ensemble is defined based on the absolute distance of its orientation from the reference orientation, categorized into one of 4 bins, equally-spaced over the range [0, 45] deg. Subjects’ choices across trials are used as the dependent variable. “Robust averaging” refers to a nonuniform weighting profile that shows higher weights for items in the ensemble with orientations close to the decision boundary (inlier bins) and lower weights for items with orientations further away (outliers bins). We used the same regression analysis to recover the weighting profile of the model predictions. Specifically, we run the model on the same stimulus ensembles used in our psychophysical experiment, computed the model’s decision probability *p*(*Ĉ*) for each trial (see below), and then performed the above regression analysis on the predicted decision probabilities.

#### Efficient ensemble coding model

In the following, we provide a brief description of the model first introduced in Ni and Stocker (2023). We refer the reader to the original article for more details. The model is composed of an efficient encoding stage and an optimal Bayesian decision stage. The core assumption of the model is that the observer has limited sensory resources/bandwidth and therefore seeks to efficiently use those coding resources for the task. More specifically, we assume that for a given overall resource limit, sensory encoding is aimed at maximizing the mutual information *I*[*s, x*] between the stimulus feature *s* and its noisy sensory representation *x* (Linsker, 1988; Wei and Stocker, 2015). This assumption imposes a constraint on encoding precision (measured as Fisher information *J* (*s*)) in terms of the stimulus distribution *p*(*s*) such that

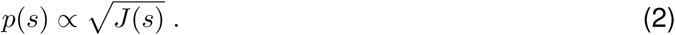

As in Ni and Stocker (2023), we assume that the prior *p*(*s*) is determined by the overall orientation distribution of the ensemble items in the experiment relative to the reference. Given our experimental design, this relative distribution is approximately Gaussian around the reference orientation (see Fig. 2b, left panel). The efficient coding constraint Eq. (2) states that more frequently occurring orientations are encoded with higher precision (i.e., higher Fisher information; Fig. 2b, middle panel). As a result, the likelihood functions of the efficient ensemble coding model are inhomogeneous, narrower for sensory representations *x* close to the reference orientation and wider and more asymmetric for representations further away (Fig. 2b, right panel). We first define a “sensory space” *s̃*, in which Fisher information is uniform. Orientation in “stimulus space” *s* is mapped to this sensory space via the cumulative of the prior distribution in the stimulus space *F* (*s*), hence

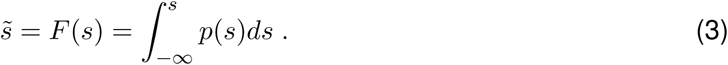

We further assume that the measurement noise and thus the likelihood function is homogeneous and symmetric in sensory space. More specifically, we assume that the noisy measurement *x* follows a von Mises distribution with mean *s̃* and a concentration parameter *κ*, thus

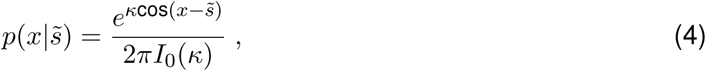

where *I*_0_ is the modified Bessel function of order zero. After defining the symmetric likelihood function in the sensory space, we can map it back to the stimulus space via the inverse mapping function *F* ^−1^(*s̃*). The resulting likelihood function *p*(*x*|*s*) in stimulus space is typically asymmetric, with long tails away from the peak of the prior distribution (see Fig. 2b, right panel).

The optimal decision process consists of performing inference over the hierarchical generative model illustrated in Fig. 2d. The generative model reflects the statistical structure of our psychophysical decision experiment. On every trial, the ensemble category *C* = {cw, ccw} is randomly chosen with equal probability *p*(*C*) = <u>^1^</u> . The chosen category determines the distribution from which the orientations of the items in the ensemble are sampled. Specifically, given each ensemble category, orientations are independently sampled from a Gaussian with mean *µ_i_*and standard deviation *σ_j_*. The generative mean *µ_i_* is randomly selected from a predefined set *µ* = {*µ*_1_*, . . . , µ_m_*}, while *σ* is fixed. This defines the probability of the feature value of each item in the ensemble in a given trial as *p*(*S*|*µ_i_, σ*) with *S* = (*s*_1_*, . . . , s*_12_) representing the 12 orientations of the items in the ensemble. Finally, the generative model considers that the observer only has access to noisy sensory representations of the ensemble stimuli *p*(*X*|*S*), where the feature values of individual stimuli are assumed to be independently but efficiently encoded according to *p*(*x_i_*|*s_i_*) as described above.

In the experiment, the observer’s task is to choose the category with highest posterior probability *p*(*C*|*X*) given the measurement array *X* = (*x*_1_*, x*_2_*, …, x_n_*). Using Bayes’ rule, we can define a decision variable *d* as the posterior ratio

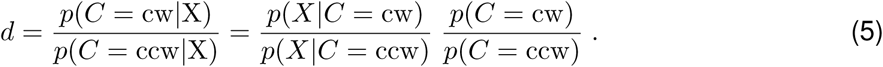

Category likelihoods *p*(*X*|*C* = cw) and *p*(*X*|*C* = ccw) are computed by summing over all possible combinations of *µ_i_* and *σ_j_* within each category. Likewise, the observer needs to marginalize over all possible orientation values *s* given each measurement *x_i_*. Therefore, the decision variable is given by

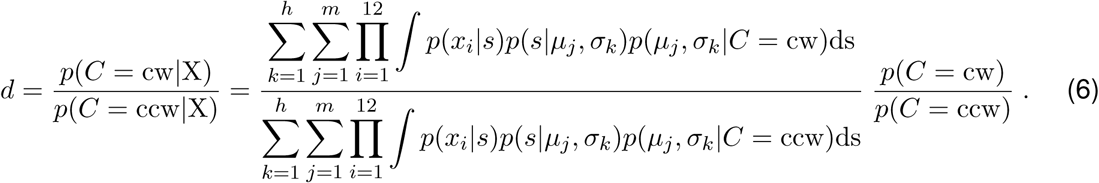

The model assumes that the observer reports a decision *Ĉ* = cw when *d >* 1, and *Ĉ* = ccw otherwise. To predict the probability of the observer’s decision on each trial, we marginalize over all possible measurements *x_i_* given each item’s orientation *s_i_*. Thus, the response probability given the stimulus ensemble *S* is

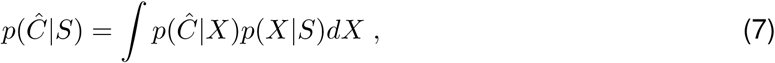

where *S* represents the orientations of the items in the ensemble as defined above and *X* corresponds to all possible, independent measurements of these orientations. We numerically perform marginalization by computing the empirical decision probability based on 4000 random samples of the measurement distributions Eq. (4) for every ensemble configuration in the experiment. Note that all model predictions shown in the paper are based on the same sets of ensemble stimuli as used in the experiment.

#### Model fits

The efficient Bayesian observer model has two free parameters:

- *κ* – parameter that determines the overall sensory encoding precision; it specifies the width of the homogeneous noise distribution in sensory space.
- *β* – parameter that determines how much efficient sensory encoding is adapted to the stimulus distribution; it specifies the relative contributions of the stimulus distribution in the experiment and a uniform distribution to the stimulus prior *p*(*s*). Efficient encoding is determined according to *p*(*s*) (Eq. 2). We model *p*(*s*) as a weighted sum of the dynamic stimulus distribution *p_e_*(*s*) relative to the decision boundary and a uniform distribution *p_u_*(*s*), with *β* and 1 − *β* being their relative contributions (see Fig. 2c).

We determined the optimal parameter values that minimized the negative log-likelihood of the models given the data using an adaptive search algorithm (Acerbi and Ma, 2017).

## Results

We first analyzed the impact of the inter-stimulus interval (ISI) on task performance and weighting profile at a coarse time-scale, grouping the trials in the ensemble-reference and reference-ensemble conditions each into two equally sized timing bins, (0, 0.45] s and (0.45, 0.9] s, respectively. Together with the simultaneous presentation condition, this resulted in five different timing conditions. Figure 3a shows the average decision accuracy across subjects (*N* = 9) in the different timing conditions. Generally, decision accuracy was higher when the ensemble mean *µ* was further away from the decision boundary. Overall decision accuracy was highest for the simultaneous condition (black) and lowest for the reference-ensemble condition (blue). Decision accuracy for the ensemble-reference condition lied in between (red). Interestingly, while performance was markedly different between the three general timing conditions (simultaneous, ensemble-reference, reference-ensemble), performance was indistinguishable between short and long ISI durations.

**Figure 3:**
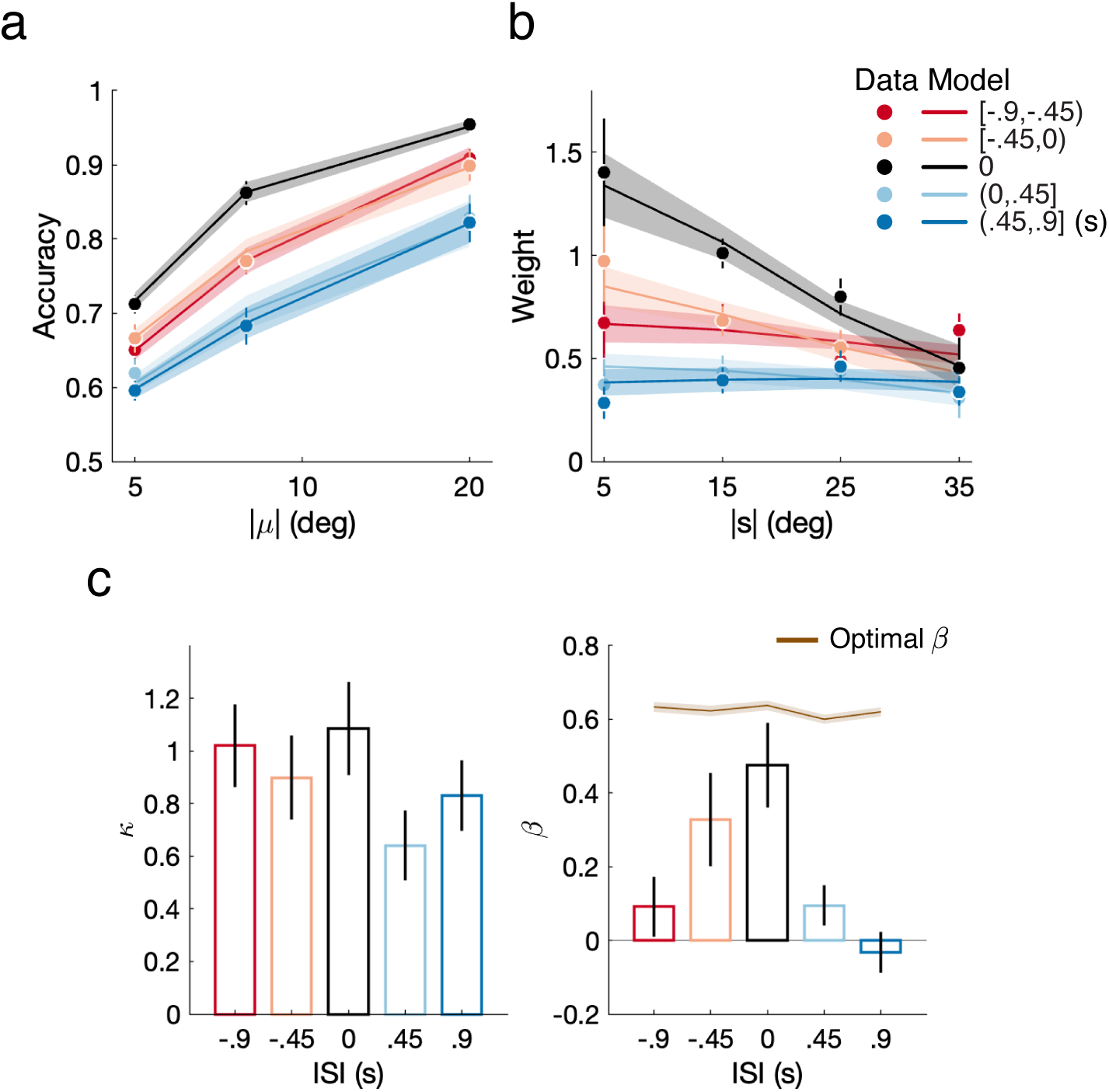
Data and model fits. (a) Decision accuracy and (b) recovered regression weights from the data (dots) and the fit model (lines) as a function of the distance of the ensemble mean from the reference orientation for different ISI conditions. Negative and positive ISI indicate the ensemble stimuli preceding (red) or succeeding (blue) the reference, respectively, with 0 s ISI representing the simultaneous presentation of the ensemble and reference stimuli (black). (c) Fit *κ* (overall encoding precision) and *β* (weight of the stimulus distribution in the prior mixture) for each of the five ISI conditions. The brown line indicates the optimal *β* that produces the highest accuracy given the fit *κ* in each condition (see Fig. 4). Shown are the combined data across all subjects. Error bars and shaded areas represent SEMs of the data and model fits, respectively. Supplementary Fig. A1: Decision accuracy, regression weights, and model fits for individual subjects’ data.

A temporal offset between the presentation of the ensemble and the reference stimuli had a substantial impact on the recovered regression weights (Fig. 3b). Robust averaging (i.e., inlier overweighing) was strongest in the simultaneous condition but markedly reduced/absent in both the ensemble-reference and reference-ensemble trial conditions. Efficient ensemble coding thus relies on simultaneously evoked sensory representations of both the ensemble and the reference stimuli. While the weighting profile was essentially flat for trials in which the reference preceded the ensemble stimulus irrespective of ISI, a weak efficient coding effect remained when the ensemble preceded the reference stimulus within a small ISI (light red). Next, we separately fit our efficient ensemble coding model to the data in the simultaneous condition and both time-bins of the ensemble-reference and reference-ensemble conditions. The model accurately captures the measured patterns of decision accuracy across all trials conditions (Fig. 3a, solid lines). Furthermore, using the same regression analysis as for the human data, the weighting profiles extracted from the model’s predicted trial-by-trial responses closely aligned with those recovered from the data (Fig. 3b, solid lines).

Fitting the model to each ISI condition separately provides insight into how the different ISI conditions affect efficient ensemble coding. The model has two parameters, the overall encoding precision *κ*, and the degree of adaptation of the encoding to the experimental ensemble distribution, *β*. Figure 3c shows the fit parameter values for the five different ISI conditions. The values indicate that the overall lowest decision accuracy in the reference-ensemble condition can be largely attributed to an overall less accurate stimulus representation (i.e., smaller *κ*). Furthermore, although their overall encoding precision is rather similar, the substantially higher decision accuracy in the simultaneous condition compared to the ensemble-reference condition is mainly due to a better adapted, efficient ensemble encoding as revealed by the significantly higher *β* value. Fits to all asynchronous conditions essentially resulted in *β* values close to zero, with the exception of the condition with a short ISI between the presentation of the ensemble and the reference, which has a mildly positive value. This corresponds well with the mostly uniform weighting profiles obtained from the regression analysis (Fig. 3b).

We run model simulations to more systematically evaluate how the two model parameters *κ* and *β* jointly affect predicted decision accuracy (Fig. 4). Increasing overall encoding precision *κ* monotonically increases decision accuracy. Similarly, sensory encoding that is better adapted to the ensemble stimulus distribution (higher *β* value) also leads to higher decision accuracy for a fixed overall encoding precision – but not for all ensembles equally. While decision accuracy improves for all ensemble stimuli as *β* increases from 0.1 to 0.5, it drops for ensembles with generative mean furthest away from the reference (|*µ*| = 20) as *β* further increases to 0.9 (Fig. 4a). At first sight, this appears counter-intuitive as more efficient encoding of the ensemble stimuli should lead to better performance. However, it makes sense when we recall that encoding is assumed to be efficient with regard to the overall (approximate Gaussian) distribution of the ensemble stimuli relative to the reference, and not the individual Gaussian distributions (centered at different values of *µ*) from which items in individual trials are drawn (see Fig. 2a). In particular, when *µ* is large, an increasing number of items in the ensemble are encoded with increasingly lower precision as *β* increases, because efficient coding reduces encoding resources at the tails of the overall distribution. As a result, improved encoding precision for ensembles with small *µ* (5 and 10 deg) is offset by decreased encoding precision for ensembles with large *µ* (20 deg). Thus, there is an optimal *β <* 1 that creates the highest overall (average) decision accuracy for a given overall encoding resource budget *κ* (Fig. 4b). Given the specific design of our decision experiment (i.e., ensemble distributions), the model predicts for every *κ* an optimal *β* that yields the highest decision accuracy. Interestingly, the *β* value from the model fit to the data in the simultaneous presentation condition closely approaches this optimum (Fig. 3c), suggesting that subjects in our experiment indeed efficiently allocated their limited resources according to the dynamic stimulus distribution. But they did so only to the degree that it improved their overall decision accuracy in the specific task.

**Figure 4:**
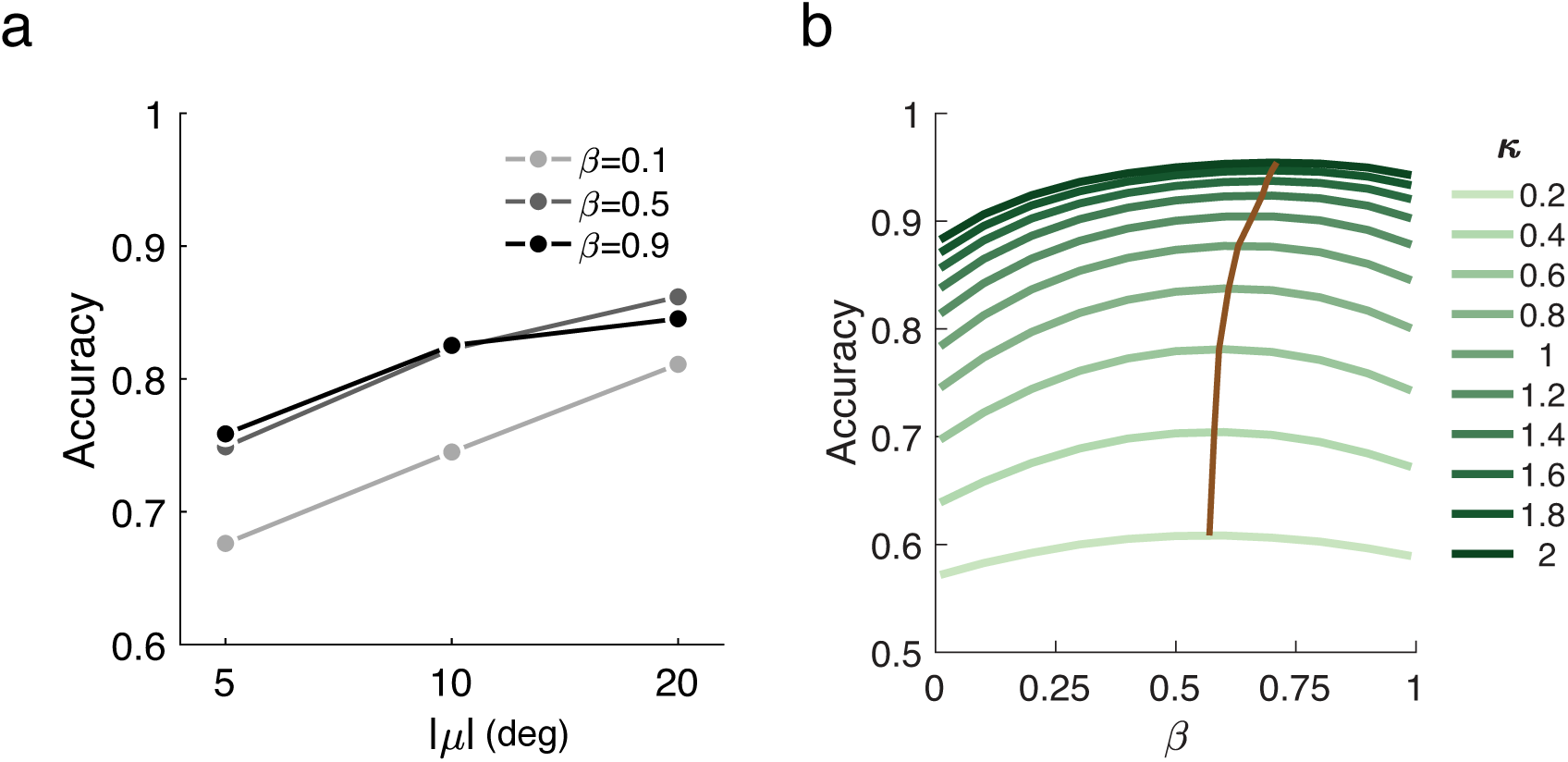
Model simulations: Decision accuracy as a function of efficient coding strength. (a) Predicted accuracy for trials sampled from different *µ* when *β* is 0.1, 0.5, and 0.9 and *κ* is fixed at 1. (b) Predicted accuracy averaged across all *µ* as a function of efficient coding strength, *β*, given different overall encoding precision *κ*. The brown lines indicate the optimal *β* values that produces the highest decision accuracy for a given overall encoding precision.

Furthermore, our analysis of the binned data suggests that the fit *β* value tends to decrease as the ISI increases in particular in the ensemble-reference condition, which is consistent with the extracted regression weight profile (Fig. 4). To examine in more detail how different ISIs affect efficient ensemble coding, we performed a sliding time-window analysis of our data. Specifically, we extracted the decision accuracy and the regression weights for trials within a 300 ms ISI timewindow sliding at 30 ms time-steps. For each time-window, we computed the difference between the average weight of the two inlier bins versus the average weight of the two outlier bins; a larger positive weight difference thus indicates stronger robust averaging (see Methods). Figures 5a,b show the resulting decision accuracies and weight differences, respectively. Decision accuracy does not significantly depend on ISI, showing a mild peak at around 350 ms and a subtle decline for longer ISIs, especially in the condition where the ensemble preceded the reference (red). Similar dynamics are also evident in the extracted weight differences. While the weight differences are essentially zero across the entire ISI range in the reference-ensemble condition (blue), they are positive early in the ensemble-reference condition before decreasing to zero due to increasing information loss during the memory period. Strikingly, neither the decision accuracy nor the weight difference showed a monotonic tendency to reach the levels observed in the simultaneous trial condition for decreasing ISIs, suggesting that the simultaneous presentation of ensemble and reference represents a unique timing condition for the decision process (Fig. 5 a and b, black dots).

**Figure 5:**
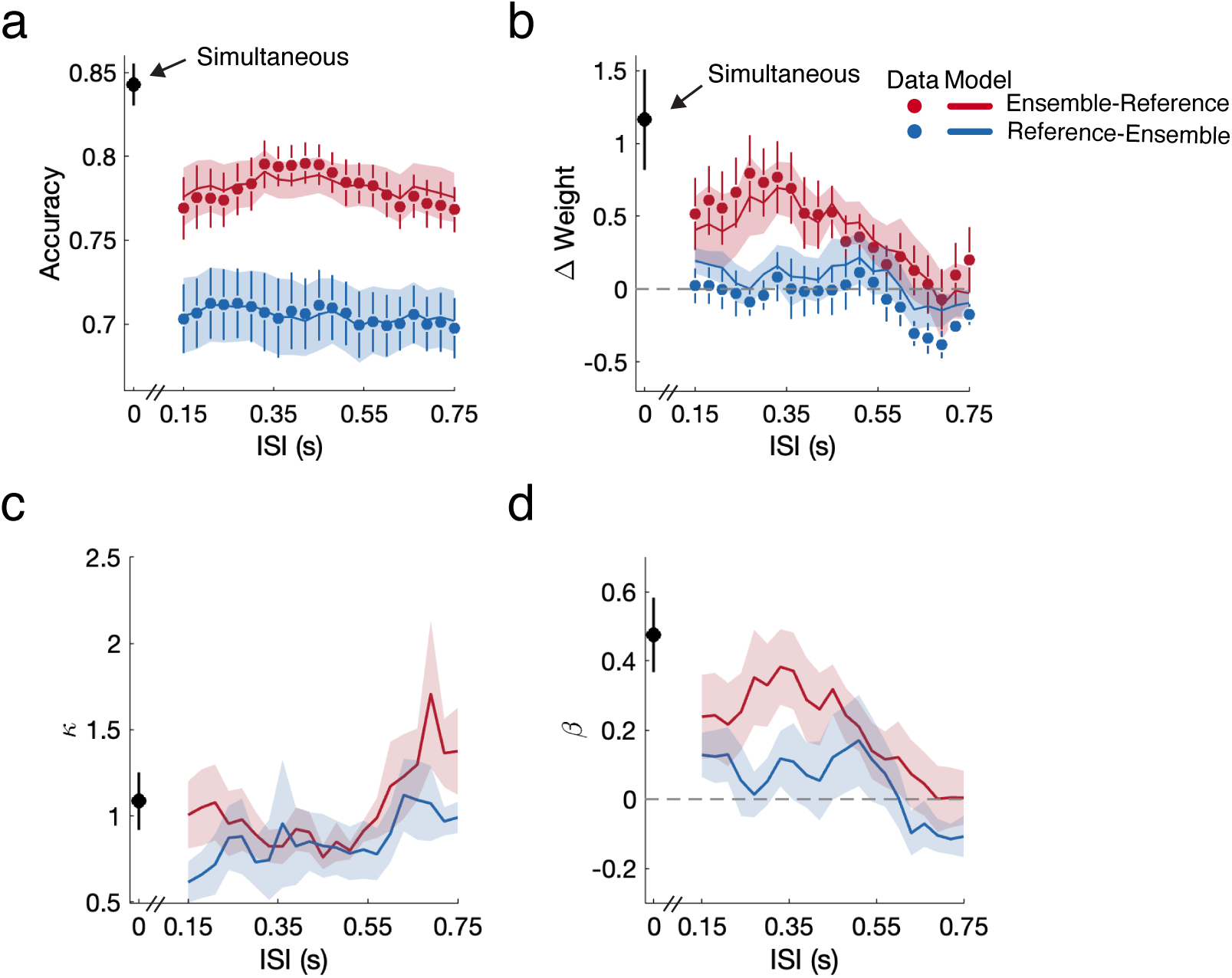
Sliding time-window analysis. (a) Overall decision accuracy of subjects (dots) and the fit model (lines) within a sliding time-window for ensemble-reference (red) and reference-ensemble (blue) conditions. (b) The observed and predicted weight difference Δ*w* between the inlier (bins 1 and 2) and outlier (bins 3 and 4) bins for each trial condition. (c) and (d) show the fit *κ* and *β* values as a function of ISI. See Methods for details.

Our model, separately fit to data of trials within each sliding ISI window, accurately tracks the changes in decision accuracy and weighting profile as a function of ISI (Fig. 5a and b, solid lines). The corresponding fit model parameter provides further insight into the dynamics of ensemble coding. First, the overall encoding resource *κ* was not markedly different between all conditions, including the simultaneous presentation condition (Fig. 5c). If anything, there is a slight tendency for higher *κ* values as ISI increases, which can be interpreted as the visual system’s attempt to compensate for the poorer performance at high ISIs with an increase in overall allocated encoding resource (e.g., through increased attention). Second, the fit parameter *β* accounts for much of the differences between the different presentation conditions, both in terms of decision accuracy and regression weights (Fig. 5d). For small ISIs in the ensemble-reference condition, *β* was substantially larger than zero, indicating that weak efficient ensemble encoding is possible if the ensemble is presented shortly before the reference. However, the fit values were significantly smaller than for the simultaneous condition with a peak at around 350 ms. Again, the trajectory of the fit *β* values between the peak and zero ISI was non-monotonic, further emphasizing that a simultaneous presentation of ensemble and reference presents a unique condition for strong efficient ensemble coding. In contrast, fit *β* values were not significantly different from zero regardless of ISI in the reference-ensemble trial condition.

## Discussion

We have previously demonstrated that the visual system can rapidly establish efficient sensory representations of stimulus ensembles relative to a dynamic reference (Ni and Stocker, 2023). This implies that efficient ensemble coding is governed by a fast resource allocation process operating at a time-scale of a few hundred ms. With the work presented here, we examined how the relative timing between the presentation of the reference and the stimulus ensemble affects this allocation process. We systematically varied the interval between the presentations of the ensemble and the reference stimuli and found that efficient ensemble coding requires the simultaneous presentation of both the ensemble and reference stimuli. Introducing a small temporal offset between the two stimuli markedly attenuates/eliminates dynamic coding reallocation.

However, some weak form of efficient ensemble coding persists when the ensemble stimulus is shown before the reference. A sliding-window analysis shows that the effect continuously decreases with increasing ISI intervals up until to about 500 ms. Surprisingly, we did not observe a similar effect when the reference presentation preceded the ensemble stimulus. We had expected that showing the reference first might even improve efficient encoding given that knowing the reference orientation ahead of time provides the system more time to allocate the encoding resources appropriately. Instead, even for very small ISIs, a preceding reference presentation immediately disrupted efficient ensemble coding, leading to a flat encoding profile. The asymmetry between the ensemble-reference and reference-ensemble conditions is also manifested in subjects’ decision accuracy and reaction times (Supplementary Fig. A2), with generally lower accuracy and longer reaction times when showing the reference first.

While our results suggest that robust averaging results from a dynamic change in sensory encoding that essentially requires neural co-activation of both the reference and the ensemble stimuli, the specific underlying mechanisms remain unknown. One candidate is a form of feature-based attention (Scolari et al., 2014; Liu, 2019). Feature-based attention is typically considered a process that enhances the responses of visual neurons whose preferred features match the attended feature value while gradually also suppressing the response of those neurons with more and more dissimilar preferences (Treue and Trujillo, 1999; Martinez-Trujillo and Treue, 2004; Maunsell and Treue, 2006). Such a feature-based gain-modulation mechanism could underlie behavioral findings showing that detection accuracy of a target stimulus gradually depends on how its feature value matches that of a pre-cued (i.e., attended) stimulus (Davis and Graham, 1981; Rossi and Paradiso, 1995; Störmer and Alvarez, 2014; Wang et al., 2015; Ho et al., 2012). However, in order to be a strong candidate mechanism for efficient ensemble coding, feature-based attention must be capable of creating gain modulation kernels that reflect learned stimulus distributions rather than a single feature value, which is currently unknown. Future neurophysiological studies will be necessary to establish this possibility.

Our results provide sufficient evidence to rule out several alternative models/explanations for robust averaging in ensemble representations. Li et al. (2017) argued that robust averaging is a way for the visual system to improve performance by making the decision more robust to late (decision) noise. If this were the case, then we would expect increased robust averaging for trials where the reference preceded the ensemble because the uncertainty in the reference orientation has increased by the time the observer has processed the (subsequent) ensemble stimulus and is ready for the decision. Our results show the opposite behavior. Likewise, Teng et al. (2021) proposed that robust averaging occurs due to a differential weighting at the decision stage, and is thus not the result of dynamic sensory encoding. If that were the case, then the relative timing of the presentation of reference and ensemble stimuli should have very little effect on the integration behavior because the same information is available at the time the decision is made (modulo some small differences in memory noise). Again, our empirical findings speak against such an interpretation. Finally, Utochkin et al. (2024) recently proposed a population response model for ensemble coding, positing that neurons with preferred feature values near the center of the stimulus distribution are over-activated because they receive stimulation from both sides of the distribution. These selective neurons with heightened activation contribute more to the ensemble average estimate, leading to an over-weighing of inlier stimuli. If this were the case, the temporal offset between the presentation of the reference and ensemble stimuli should have minimal impact on the weighting kernel, a prediction that is also contradicted by our data.

The proposed efficient ensemble coding model provides a normative explanation for robust averaging in ensemble perception. Robust averaging improves decision accuracy for a given, limited encoding bandwidth. The performance improvement occurs because the visual system adaptively allocates its encoding bandwidth/resources to process a constrained range of ensemble stimulus features based on their statistics relative to the reference. Consequently, although the total encoding resource remains constant, the representation of a stimulus ensemble is more accurate under efficient compared to uniform encoding. With only two free parameters, our model can accurately describe subjects’ behavior across all tested conditions, both in terms of their decision accuracy as well as their ensemble weighting profiles. It provides a parsimonious, normative framework for understanding ensemble encoding and decision behavior. As such, it naturally does not explicitly express all the specific cognitive processes that are likely involved in performing the task, such as working memory decay or decision noise. Thus a careful interpretation of the fit model parameters is required. For example, the model predicts slightly lower encoding precision in the ensemble-reference and reference-ensemble conditions compared to the simultaneous condition. This likely does not reflect an actual difference in encoding precision of the stimuli, but rather subsumes the precision loss during the additional memory phase in the non-simultaneous conditions. Refining the model to incorporate such additional processes (e.g., memory noise and decision noise) will help us to further probe and, ultimately, uncover the underlying mechanisms that govern efficient ensemble coding.

## Conclusion

We examined the temporal aspect of efficient ensemble coding in a discrimination task by systematically varying the temporal offset between the presentation of the stimulus ensemble and the reference stimulus. We show that a simultaneous presentation of both the ensemble and reference stimuli is critical for the visual system to establish rapid changes in the sensory encoding precision of ensemble stimuli. Our findings provide strong evidence that efficient coding can be near-instantly established.

## Appendix

**Figure A1:**
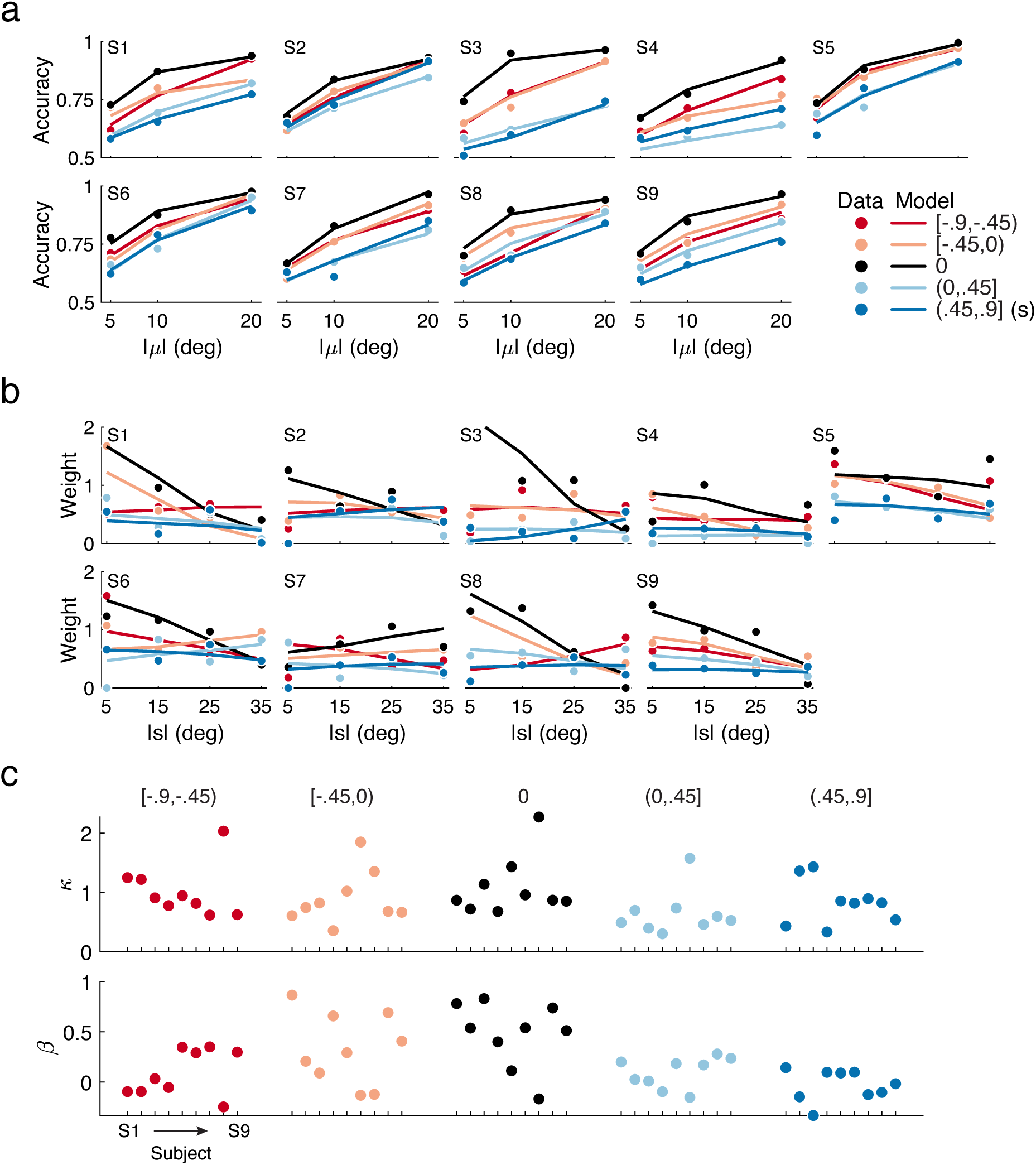
Data and model fits for individual subjects (*N* =9). (a) Decision accuracy and (b) recovered regression weights for data (filled dots) and model fits (solid lines). Different colored lines indicate condition wherein the inter-stimulus interval (ISI) was within [-0.9, -0.45] (dark red), [-.45 0] (light red), [0 0.45] (light blue), [0.45 0.9] (dark blue), and 0 (black) s, respectively. Negative ISI values indicate the conditions where ensemble stimuli preceded the reference stimulus, positive values correspond to the reference preceding the ensemble. 0 s ISI represents trials with simultaneous presentation of the ensemble and reference stimuli. (c) Fit parameters *κ* (top) and *β* (bottom) for each subject (dots), displayed in the same order from Subject 1 (S1) to Subject 9 (S9) for the five ISI conditions (color-coded).

**Figure A2:**
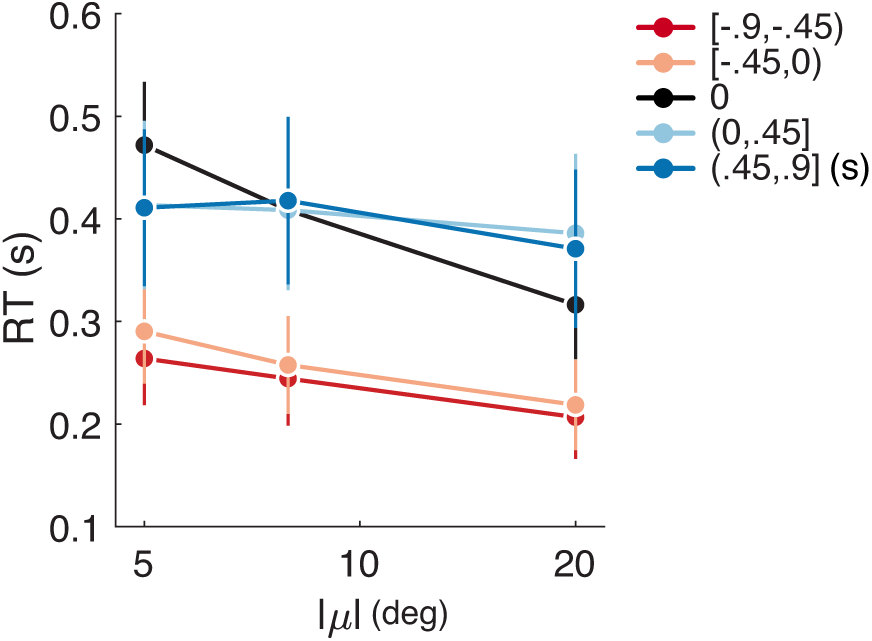
Average reaction time (RT) across nine subjects as a function of the ensemble generative mean for each of the five ISI conditions.

